# Cyto-architecture constrains a photoactivation induced tubulin gradient in the syncytial *Drosophila* embryo

**DOI:** 10.1101/520031

**Authors:** Sameer Thukral, Bivash Kaity, Bipasha Dey, Swati Sharma, Amitabha Nandi, Mithun K. Mitra, Richa Rikhy

## Abstract

*Drosophila* embryogenesis begins with nuclear division in a common cytoplasm forming a syncytial cell. Morphogen gradient molecules spread across nucleo-cytoplasmic domains to pattern the body axis of the syncytial embryo. The diffusion of molecules across the syncytial nucleo-cytoplasmic domains is potentially constrained by association with the components of cellular architecture, however the extent of restriction has not been examined so far. Here we use photoactivation (PA) to generate a source of cytoplasmic or cytoskeletal molecules in order to monitor the kinetics of their spread in the syncytial *Drosophila* embryo. Photoactivated PA-GFP and PA-GFP-Tubulin within a fixed anterior area diffused along the antero-posterior axis. These molecules were enriched in cortical cytoplasm above the yolk-filled center suggesting that the cortical cytoplasm is phase separated from the yolk-filled center. The length scales of diffusion were extracted using exponential fits under steady state assumptions. PA-GFP spread to greater distance as compared to PA-GFP-Tubulin. Both gradients were steeper and more restricted when generated in the center of the embryo probably due to a higher density of nucleo-cytoplasmic domains. The length scale of diffusion for PA-GFP-Tubulin gradient increased in mutant embryos containing short plasma membrane furrows and disrupted tubulin cytoskeleton. The PA-GFP gradient shape was unaffected by cyto-architecture perturbation. Taken together, these data show that PA-GFP-Tubulin gradient is largely restricted by its incorporation in the microtubule network and intact plasma membrane furrows. This photoactivation based analysis of protein spread across allows for interpretation of the dependence of gradient formation on the syncytial cyto-architecture.

## Introduction

Insect embryos initiate their development in a large syncytial cell where multiple nuclei undergo nuclear divisions in a common cytoplasm without forming complete cells. The cytoplasm is thought to mix uniformly in the syncytial cells. However, syncytial *Drosophila* embryos have distinct domains of gene expression patterns in nuclei despite being in this common cytoplasm (Shvartsman *et al*., 2008). Several tissues in different organisms, for example, plant endosperm cells, animal muscle cells and fungal hyphae, also contain syncytial cells. Syncytial nuclei in fungi maintain distinct cell cycle stages (Anderson *et al*., 2013; Dundon *et al*., 2016). The spatially separated daughter nuclei in these fungi continue to proceed through the cell cycle synchronously by maintaining a similar concentration of cell cycle components (Lee *et al*., 2013). Syncytial nuclei in muscle cells have a differential expression of mRNAs as compared to their neighbors (Pavlath *et al*., 1989). These studies indicate that several components of the cytoplasm have local function and are likely to be generated and sequestered in the vicinity of the syncytial nuclei. It is of interest to understand the cellular mechanisms that regulate compartmentalized distribution of molecules despite being in a common cytoplasm.

The syncytial embryos of *Drosophila* provide a tractable system to decipher the extent to which different cellular components are shared across nucleo-cytoplasmic domains. *Drosophila* embryogenesis begins with 9 nuclear division cycles deep within the embryo during the preblastoderm stage. Nuclei along with centrosomes migrate to the cortex in nuclear cycle 10 and the nuclear division cycles 11-14 occur beneath the cortex in the syncytial blastoderm embryo (Foe and Alberts, 1983; Karr, 1986; Warn, 1986; Foe,Odell and Edgar, 1993; Sullivan and Theurkauf, 1995). Each interphase nucleus of the syncytial blastoderm embryo is surrounded by apical centrosomes and a microtubule array in an inverted basket conformation. Astral microtubules reach out from the centrosomes towards the cortex and overlap with the astral microtubules originating from neighbouring nuclei (Cao *et al*., 2010). F-actin is present in caps above the nuclei and centrosomes. Lipid droplets and yolk are enriched at the bottom of the basket (Kuhn *et al*., 2015; Mavrakis *et al*., 2009a; Schmidt and Grosshans, 2018; Welte, 2015). Each nucleo-cytoplasmic domain in the blastoderm embryo is associated with organelles such as the endoplasmic reticulum, Golgi complex and mitochondria (Frescas *et al*., 2006; Mavrakis *et al*., 2009b, Chowdhary *et al*., 2017). The microtubule and the actin cytoskeleton remodel during prophase and metaphase of the syncytial division cycle. The centrosomes move laterally during prophase and give rise to spindles during metaphase. Actin is enriched along the cortex at the extending plasma membrane furrows (Foe, Odell and Edgar, 1993). The short furrows present in interphase between adjacent nuclei extend deeper between spindles in metaphase. Molecules in the plasma membrane, ER, Golgi complex and mitochondria have limited exchange between adjacent nucleo-cytoplasmic domains in the syncytial *Drosophila* embryo (Frescas *et al*., 2006; Mavrakis *et al*., 2009b, Chowdhary *et* al.,2017).

Analysis of exchange of molecules in the cytoplasm or cytoskeleton across the syncytial nucleo-cytoplasmic domains remains to be documented in a systematic manner, though several studies have probed various cytoplasmic properties. Fluorescent dextran of various sizes when injected in the cytoplasm of the syncytial blastoderm embryo has been used to estimate the rate of cytoplasmic diffusion in the embryo (Gregor *et al*., 2005). Micro-rheology based measurements of cytoplasmic viscosity have found that cytoplasmic viscosity is three times higher that of water in the region between nuclei and yolk of the syncytial *Drosophila* embryo. In addition, microtubules, but not actin contribute to the observed viscosity (Wessel *et al*., 2015).

Morphogen gradient formation in the syncytial *Drosophila* can be used as a paradigm to estimate properties of the embryo cytoplasm. Bicoid forms a gradient in the antero-posterior axis, patterning the head of the embryo (Gregor *et al*., 2007). The Bicoid gradient is present primarily in the cortical region of the embryo (Cai *et al*., 2017). The dorso-ventral gradient formed by Dorsal is compartmentalized to each nucleo-cytoplasmic domain (DeLotto *et al*., 2007) and modelling studies show that plasma membrane furrows could restrict Dorsal gradient spread (Daniels *et al*., 2012). The Dorsal gradient formation on the ventral side depends on specific binding partners on the ventral side (Carrell *et al*., 2017). These studies together imply that the syncytial blastoderm cortex shows gradients whose properties depend upon sequestration due to interaction with other cytoplasmic components or the syncytial cyto-architecture.

In this study, we attempt to elucidate the extent of gradient spread across nucleo-cytoplasmic domains of the syncytial *Drosophila* embryo using a comparison between cytoplasmic PA-GFP and PA-GFP-Tubulin. Fluorescently labelled tubulin incorporates well in the microtubule network and is also present in the cytoplasm. We use photoactivation to generate a fixed population of PA-GFP or PA-GFP-Tubulin and find that both diffuse in the cortical region as compared to the yolk filled central region of the syncytial blastoderm embryo. The gradient of PA-GFP-Tubulin is more restricted as compared to PA-GFP in the antero-posterior axis. PA-GFP and PA-GFP-Tubulin have a decreased spread when generated in the middle of the embryo as compared to the anterior. The PA-GFP-Tubulin gradient diffuses to a greater distance in mutants showing a loss of plasma membrane furrows and disruption of the microtubule network. The PA-GFP gradient is not affected in these mutants. Our study provides a framework for assessing the regulation of gradient formation by its interaction with the syncytial cytoarchitecture components and has implications on the spread of morphogen gradients across different paradigms.

## Results

### Cytoplasmic GFP and mCherry-Tubulin are enriched cortically in the syncytial division cycles in the *Drosophila* embryo

The syncytial *Drosophila* blastoderm embryo has a characteristic arrangement of microtubules around each nucleus. Microtubules emanate from the apical centrioles and spread vertically covering the nuclei in an inverted basket like arrangement (Karr, 1986; Sullivan and Theurkauf, 1995). In order to test the extent of spread of molecules in the cytoplasm we imaged embryos expressing GFP ubiquitously under the control of the *ubiquitin* promoter. GFP is expected to be present primarily in the cytoplasm and is not known to interact with any cytoplasmic components (Verkman, 1999). We compared the expression of cytoplasmic GFP to fluorescently labelled tubulin as it would partition into the cytoplasm and also incorporate into the microtubule cytoskeleton. For this we imaged live embryos expressing fluorescently tagged alpha-Tubulin (UASp-mCherry-Tubulin) (Rusan and Peifer, 2007) with *mat-Gal4-vp16* (mat-Gal4). We found that cytoplasmic GFP was enriched cortically and accumulated inside the cortical nuclei (Figure 1A). Accumulation of GFP occurs passively inside the nucleus as a result of its small size which allows it to pass through the nuclear pore complex (Ruiwen Wang, 2007). The fluorescence intensity of cytoplasmic GFP progressively increased near the cortex as syncytial division cycles progressed but remained above the yolk filled region (Figure 1B, Movie S1). We noticed GFP fluorescence dropped to approximately 30% between 32 to 36μm in syncytial cycle 14 (Figure 1E). mCherry-Tubulin was enriched on apical centrioles, in microtubules spreading vertically from the cortex and in the cytoplasm in the syncytial division cycles (Figure 1C, Movie S2). mCherry-Tubulin also showed progressive accumulation of fluorescence signal near the cortex as the syncytial cycles progressed (Figure 1D). mCherry-Tubulin fluorescence dropped to 30% between 25 to 27μm beneath the cortex in syncytial cycle 14 (Figure 1F). Thus cytoplasmic GFP and mCherry-Tubulin were concentrated near the cortex and further enriched during the progression of the nuclear cycles. In addition, they were present in a separate cortical layer of cytoplasm on top of and distinct from the inner yolk-filled region of the embryo.

**Figure 1:**
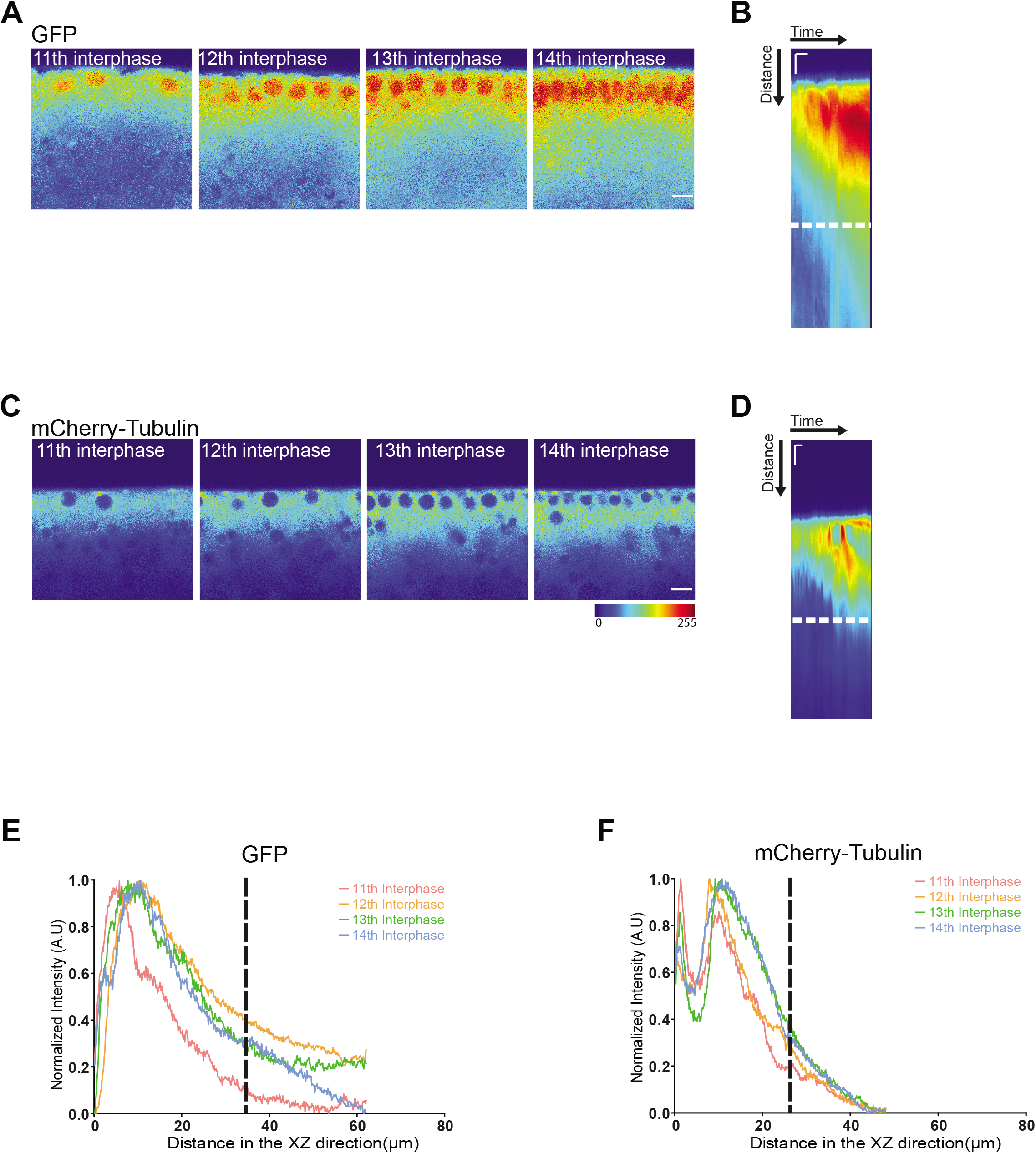
Cytoplasmic GFP and mCherry-Tubulin are enriched cortically in the syncytial division cycles. A-D: Characterization of cortical spread of GFP and mCherry-Tubulin in the syncytial division cycles. Images are shown from different cycles (NC11,12,13,14) of embryos ubiquitously expressing GFP (A) or maternally expressing mCherry-Tubulin (similar trends were observed for n=3 movies) (C). Kymographs show cortical enrichment of fluorescent signal for GFP (B) and mCherry-Tubulin (D) over time. Scale bar=5μm, 600s. E-F: Quantification of cortical enrichment of fluorescent signal in cytoplasmic GFP and mCherry-Tubulin. Graph shows normalized intensity profile for GFP (E) and mCherry-Tubulin (F) obtained from a line drawn from the cortical region towards the centre of the embryo. The dashed line shows a point at which the intensity drops to 30%. Note that the signal remains above the region containing the dark yolk filled vesicles. The images are shown in a 16 color intensity rainbow where Blue represents the lowest intensity and red represents the highest intensity. Scale bar= 10μm

### Photoactivation generates a source of PA-GFP and PA-GFP-Tubulin at the anterior that forms a cortical gradient along the antero-posterior axis

Labelled tubulin had a cytoplasmic and a microtubule bound fraction, in contrast to GFP, which had a cytoplasmic fraction in the *Drosophila* syncytial blastoderm embryo. This gave us an opportunity to assess the diffusion of these two proteins in the cytoplasm across nucleo-cytoplasmic domains. Computational simulations have predicted that binding to microtubule network and movement on motors is sufficient for partitioning the cytoplasm, in the absence of membrane boundaries in the syncytial blastoderm embryo (Chen *et al*., 2012). We therefore asked whether tubulin which partitions partially into microtubules could be more restricted as compared to GFP in the syncytial blastoderm embryo.

Photoactivation of cytoplasmic and cytoskeletal proteins has been used to generate a local source of protein molecules for monitoring their directional spread in axons (Gauthier-Kemper *et al*., 2012; Gura Sadovsky *et al*., 2017). In order to differentially test the spread of cytoplasmic and cytoskeletal proteins in the syncytial blastoderm embryo, we used photoactivation to create a local source of fluorescent PA-GFP or PA-GFP-Tubulin at different locations of the embryo (Figure 2A). Unlike morphogens such as Dorsal and Bicoid, GFP and tubulin are not differentially distributed in the syncytial embryo. PA-GFP and PA-GFP-alpha-Tubulin84B (PA-GFP-Tubulin) were expressed individually in embryos by crossing the transgenic flies to mat-Gal4. A fixed area was continuously photoactivated to form fluorescent PA-GFP/PA-GFP-Tubulin, thus creating a local source of PA-GFP/PA-GFP-Tubulin at the anterior pole of the embryo (Figure 2B,D, Movie S3,4). The movies of PA-GFP photoactivation also showed the presence of a strong autofluorescent signal at the base of the cortex comprising of yolk (Movie S3). The movies of PA-GFP-Tubulin showed an increase in fluorescence in the cytoplasm and PA-GFP-Tubulin was also incorporated in microtubules in interphase and in metaphase spindles (Movie S4). Both PA-GFP and PA-GFP-Tubulin increased in concentration by diffusion away from the source across the syncytial division cycles. A kymograph obtained at the source of photoactivation showed a distinct increase in amount of photoactivated molecules over time (Figure 2C,E). The kymograph also showed that the fluorescent signal was enriched near the cortex and did not enter the central yolk filled region of the embryo. An analysis of the directionality of spread showed that both molecules spread to a greater distance cortically along the antero-posterior axis (XY) as compared to the depth within the embryo (XZ) (Figure 2F,G). The cytoplasm of syncytial *Drosophila* blastoderm embryo has a biphasic distribution with cortical nucleo-cytoplasmic domains present above a barrier comprising of yolk and other unknown components (Foe and Alberts, 1983; Wessel *et al*., 2015). This organization possibly allows for greater spread along the cortex in the antero-posterior axis as compared to the centre.

**Figure 2:**
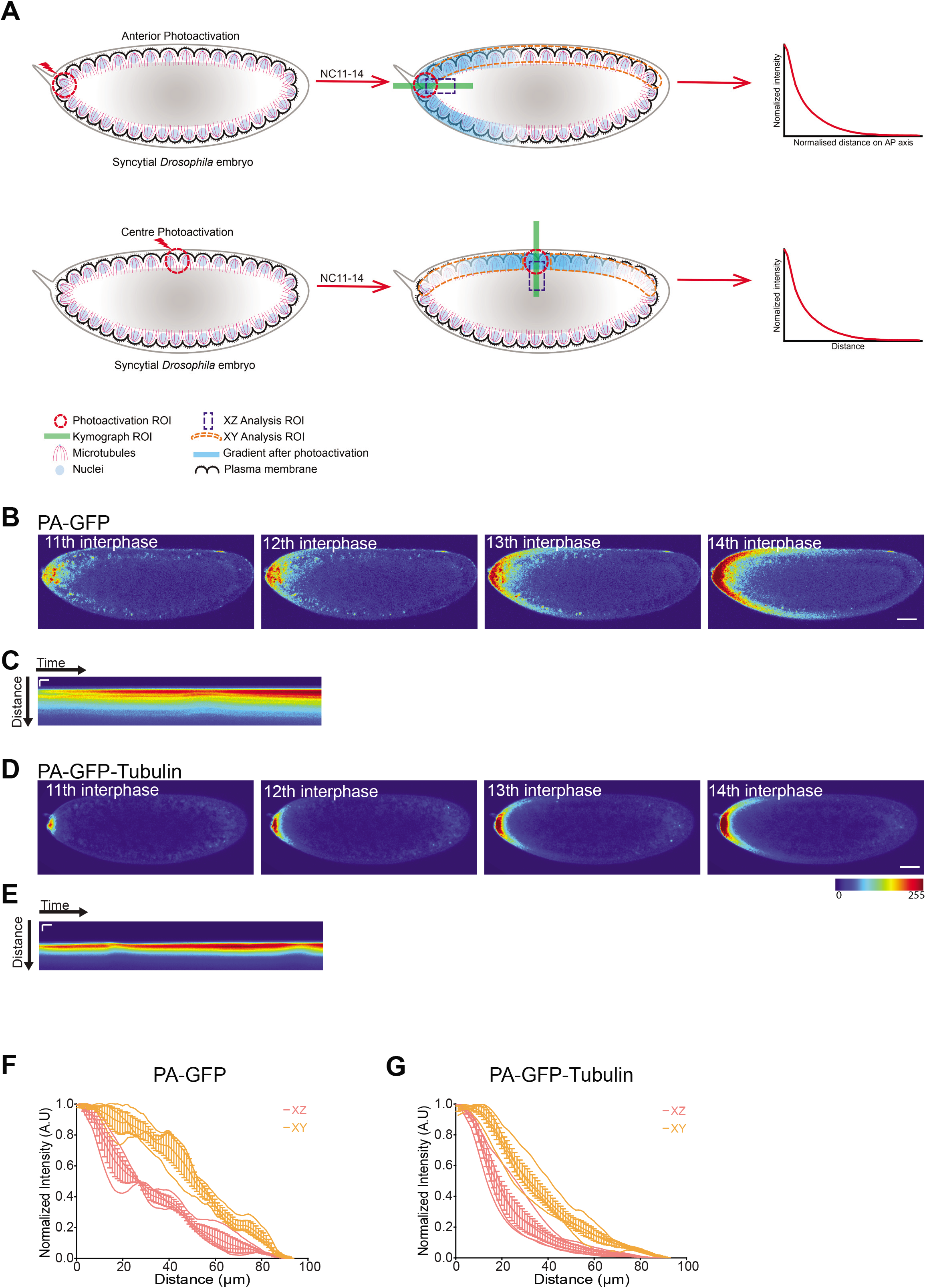
Anteriorly photoactivated PA-GFP and PA-GFP-Tubulin produces a cortical gradient. A. The photoactivation method to create an ectopic source of PA-GFP and PA-GFP-Tubulin. Photoactivation was carried out in a fixed area (373μm^2^) in the anterior or the center of the syncytial embryo. A kymograph monitoring the increase in signal was drawn at the source (green bar). A cortical region was drawn to estimate the change in intensity in the antero-posterior axis (orange). The exponential function was fit to estimate the length scale of spread for the gradients. B-E. Anteriorly photoactivated PA-GFP and PA-GFP-Tubulin forms a gradient. Images for NC11,12,13,14 of embryos from expressing PA-GFP (B) or PA-GFP-Tubulin (D) are shown after photoactivation at the anterior pole. Kymograph shows increase in cortical fluorescence over time in PA-GFP (C) and PA-GFP-Tubulin (E) expressing embryo. Scale bar=50μm,60s. F-G. PA-GFP and PA-GFP-Tubulin spreads preferentially at the cortex. Graph quantifying the extent of spread of photoactivated protein fluorescence in the planar or antero-posterior XY axis vs depth or XZ direction for PA-GFP (F) and PA-GFP-Tubulin (G) with a line drawn across either XY or XZ direction from the activated region. The raw data is in a lighter color and the averaged data is in a darker color, error bars represent standard error on means (n=3 embryos for PA-GFP-Tubulin and PA-GFP each). The images are shown in a 16 color intensity rainbow where Blue represents the lowest intensity and red represents the highest intensity. Scale bar= 50μm.

### Anteriorly photoactivated PA-GFP and PA-GFP-Tubulin shows an exponential gradient that is steeper for PA-GFP-Tubulin

We next attempted to quantify the gradients obtained in the antero-posterior axis on photoactivation of PA-GFP and PA-GFP-Tubulin anteriorly. We found that the photoactivated probes spread further as the syncytial cycles progress (Figure 3A,B). The fluorescence intensity of PA-GFP and PA-GFP-Tubulin increased with time at different locations in the embryo (Figure 3C,D). The concentration of PA-GFP and PA-GFP-Tubulin when measured at 11μm from the photoactivation source, increased with time and reached saturation. The time taken to reach steady state increased as we moved away from the photoactivation source. Temporal evolution of fluorescence at x=38μm approached a steady state at later time points. In contrast, for x=165μm, the concentration did not reach a steady value (Figure 3C,D). This is also apparent from the temporal evolution of the rate of change of the concentration at these different locations (Figure 3E,F). We used the steady state concentration profile and extracted the characteristic length scales by fitting it to an exponential decay equation (Figure 3G,H). PA-GFP and PA-GFP-Tubulin formed gradients of distinct length scales after activation at the anterior pole (Figure 3I). We found that the length scale for PA-GFP (186+25μm) was significantly higher than PA-GFP-Tubulin (70+4μm) (Figure 3J). The estimated diffusion coefficient for PA-GFP was 44.25μm^2^/s and PA-GFP-Tubulin was 20.87μm^2^/s (refer to Materials and Methods). This is likely to be because PA-GFP-Tubulin, in addition to being present in the cytoplasm is also engaged in forming the microtubule cytoskeleton and this turnover makes it less available to diffuse as compared to PA-GFP alone.

**Figure 3:**
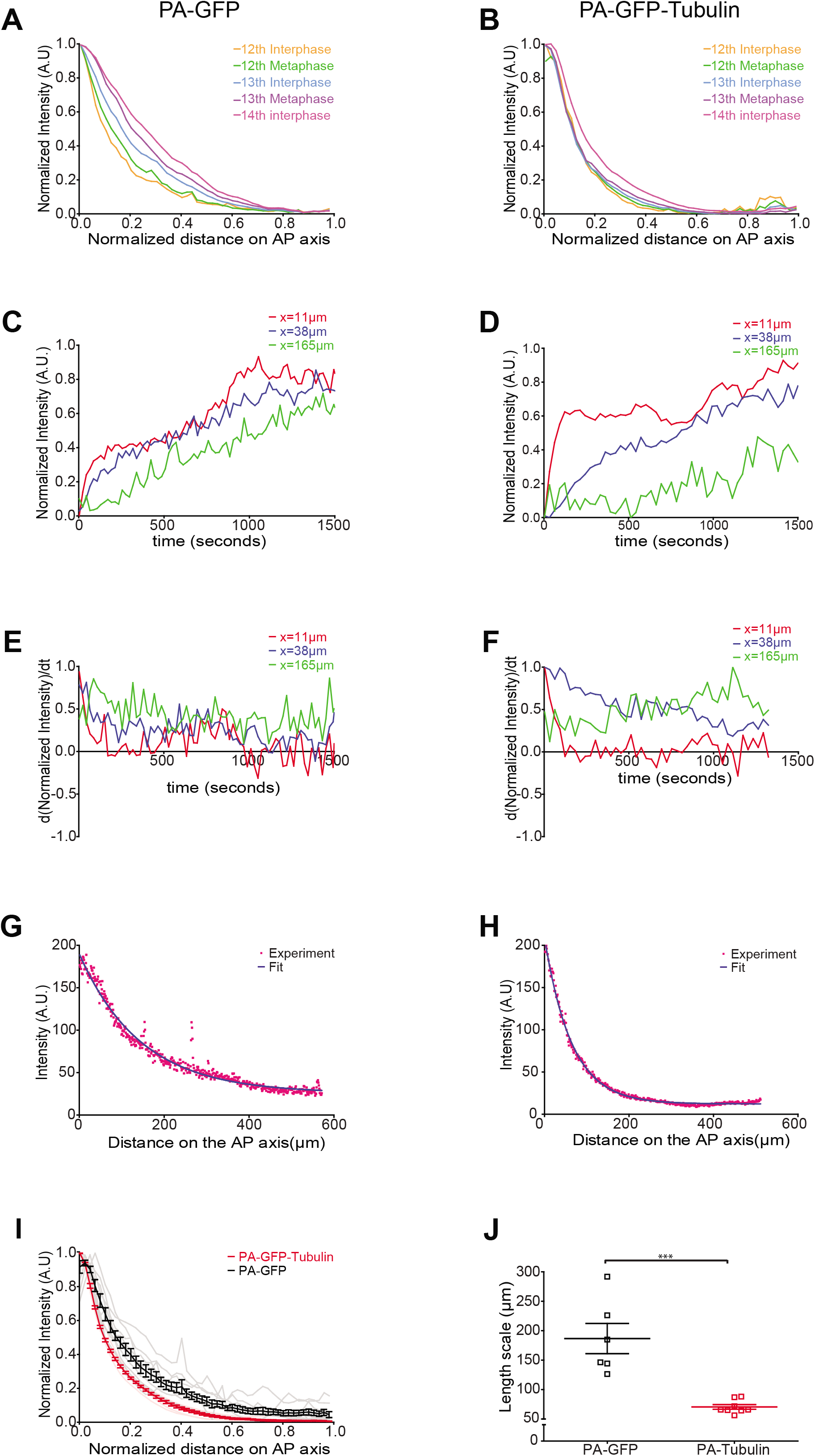
Anteriorly photoactivated PA-GFP and PA-GFP-Tubulin forms an exponential gradient with PA-GFP-Tubulin being more restricted as compared to PA-GFP. A-B. Quantification of the photoactivated signal across nuclear cycles. Graph shows intensity for PA-GFP (A) and PA-GFP-Tubulin (B) for one embryo with a line drawn across the cortical region in the syncytial nuclear cycles. Similar profiles were observed in multiple embryos (n=3 for each). C-D. PA-GFP and PA-GFP-Tubulin increases in concentration over time. The graph depicts increase in PA-GFP (C) or PA-GFP-Tubulin fluorescence intensity over time as measured at different locations (11, 38, 165μm from the source of photoactivation at the anterior). E-F. Graph shows the rate of change in concentration of photoactivated PA-GFP (E) and PA-GFP-Tubulin (F) to assess if the steady state has reached. Each plot is a derivative of the corresponding plot in C,D. G-H. Anteriorly photoactivated PA-GFP and PA-GFP-Tubulin shows an exponential gradient. Raw experimental values (red) were fit to an exponential function (blue) for each probe. I. Quantification of intensity profile of photoactivated probe measured at the end of the experiment for PA-GFP and PA-GFP-Tubulin. Graph shows raw data in a lighter color and averaged data in a darker color, error bars represent standard error on means (n=3 embryos each for PA-GFP and PA-GFP-Tubulin). J. Scatter plot of length scales extracted after fitting an exponential decay function to the intensity profiles seen in I (n=6 regions drawn in 3 embryos for PA-GFP, 8,4 for PA-GFP-Tubulin, Two tailed Mann-Whitney non-parametric test with p value=0.0007).

### Photoactivation of the cytoplasmic PA-GFP and PA-GFP-Tubulin in the middle of the *Drosophila* embryo results in a gradient with a smaller length scale as compared to the anterior activation

The syncytial *Drosophila* embryo has three domains containing distinct patterns of density of nuclei and packing (Blankenship and Wieschaus, 2001; Rupprecht *et al*., 2017). The domains show different speeds of furrow extension during cellularization. The anterior and posterior domain contain nuclei at a lower density as compared to the middle domain and the cells formed have a shorter plasma membrane furrows as compared to the middle domain in cellularization. This difference in architecture across the antero-posterior axis is regulated by the patterning molecules Bicoid, Nanos and Torso (Blankenship and Wieschaus, 2001). This difference in the density of nucleo-cytoplasmic domains prompted a comparison of the extent of gradient spread, when it originates in the middle of the embryo versus when it originates in the anterior domain (Figure 2A).

We tested if there was a difference in kinetics of gradient formation when photoactivation was carried out in the middle (Figure 4) of the embryo as compared to the anterior (Figure 2,3). For this we photoactivated PA-GFP and PA-GFP-Tubulin containing embryos in a fixed region in the middle of the embryo (Figure 2A, 4A,C). Photoactivation produced a cortical gradient with a progressive increase in gradient spread across the syncytial division cycles (Figure 4B,D-F, Movie S5,6). The gradient of PA-GFP and PA-GFP-Tubulin spread to a greater extent in the antero-posterior axis (XY) as compared to the depth of the embryo (XZ), away from the region of photoactivation (Figure 4G,H).

**Figure 4:**
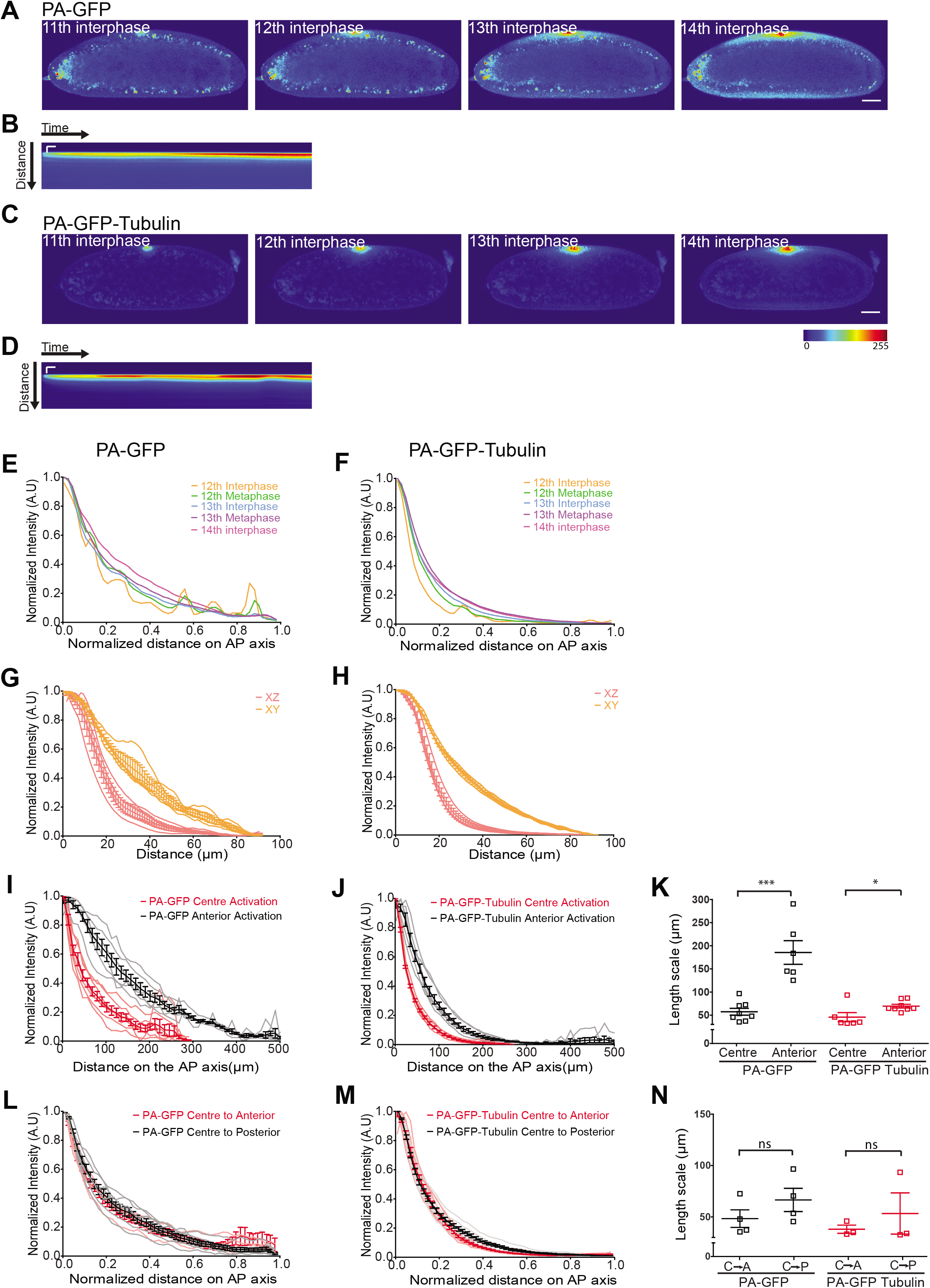
Photoactivation of the cytoplasmic PA-GFP and PA-GFP-Tubulin in the middle of the *Drosophila* embryo. A-D. Monitoring gradient of center photoactivated PA-GFP and PA-GFP-Tubulin. Images from NC11,12,13,14 expressing PA-GFP (A) or PA-GFP-Tubulin (C) and photoactivated at the centre of the embryo are shown. Kymograph shows increase in cortical fluorescence with time for PA-GFP (B) and PA-GFP-Tubulin (D) embryo. Scale bar=50μm, 60s. E-F. Quantification of evolution of photoactivated signal in syncytial nuclear cycles. The graph depicts the fluorescence intensity for PA-GFP (E) and PA-GFP-Tubulin (F) from one embryo for a line drawn from the source along the antero-posterior axis. Similar profiles were observed in multiple embryos (n=3 for each). G-H. Quantification of photoactivated protein in XY vs XZ direction for PA-GFP (G) and PA-GFP-Tubulin (H). Graph shows intensity profile of a line drawn in the XY or XZ direction from the activated region. The raw data is shown in a lighter color and the averaged data is shown in a darker color, error bars represent standard error on means (n=3 embryos for PA-GFP-Tubulin and PA-GFP each). I-J. Graphs comparing the intensity profile obtained upon photoactivation at the anterior pole versus the centre of the embryo for PA-GFP (I) and PA-GFP-Tubulin (J) (n=3 embryos for PA-GFP and PA-GFP-Tubulin each). K. Scatter plot of the length scales extracted after fitting an exponential decay function to the intensity profiles seen in I,J. The values of length scales for PA-GFP and PA-GFP-Tubulin for anterior photoactivation are repeated from Figure 3J. (n=8,4 for PA-GFP center activation, 6,3 for PA-GFP activated anteriorly, 6,3 for PA-GFP-Tubulin center activation and 8,4 for PA-GFP-Tubulin anterior activation. Two tailed Mann-Whitney non-parametric test with p value=0.0007 for PA-GFP and 0.04 for PA-GFP-Tubulin). L-M. Graphs comparing the intensity profiles obtained upon photoactivation at the centre of the embryo for analysis of directionality of spread. Fluorescence intensity is obtained from a line drawn from the centre activated region towards anterior or posterior pole for PA-GFP (I) and PA-GFP-Tubulin (J). The raw data is shown in a lighter color and the averaged data is shown in a darker color, error bars represent standard error on means (n=3 embryos for PA-GFP-Tubulin and PA-GFP each). N. Scatter plot of the length scales extracted after fitting an exponential decay function to the intensity profiles seen in L,M. (n=4,4 for PA-GFP center activation, center to anterior subset from K, 4,4 for PA-GFP center activation, centre to posterior subset from K, 3,3 for PA-GFP-Tubulin center activation, centre to anterior subset from K and 3,3 for PA-GFP-Tubulin center activation, centre to posterior subset from K. Two tailed Mann-Whitney non-parametric test with p value=0.34 for PA-GFP and 1 for PA-GFP-Tubulin). The images are shown in a 16 color intensity rainbow where Blue represents the lowest intensity and red represents the highest intensity. Scale bar= 50μm.

Length scale values were extracted by fitting an exponential equation and it was found that the extent of spread for both the probes was lower (PA-GFP 57.4+7μm, PA-GFP-Tubulin 45.6+10μm) than that observed when photoactivation was performed anteriorly (Figure 4I-K). We further analysed if there was any difference in the gradient formation from the centre towards the anterior versus centre towards the posterior pole (Figure 4L,M). We found the length scales of the gradients did not differ in either direction (Figure 4N). These analyses show that the gradient spreads uniformly across the syncytial nucleo-cytoplasmic domains towards the anterior pole and the posterior pole of the *Drosophila* embryo, negating the presence of any cytoplasmic flows or currents. In summary, photoactivated molecules generated in middle spread to a smaller distance as compared to when they were generated at the anterior pole.

### Anteriorly photoactivated PA-GFP-Tubulin gradient length scale increases in embryos containing an overexpression of RhoGEF2 on loss of pseudocleavage furrows

The gradients produced by PA-GFP and PA-GFP-Tubulin provided a framework to test the role of syncytial cytoarchitecture in regulating their diffusion. Each cortical nucleus in the syncytial blastoderm embryo of *Drosophila* contains a small ingression of the plasma membrane around it. Astral microtubules support ectopic furrows (Barmchi *et al*., 2005; Cao *et al*., 2008; Crest *et al*., 2012). The plasma membrane furrows ingress deeper in metaphase to form pseudocleavage furrows (Schmidt and Grosshans, 2018). To test the role of furrows in regulation of gradient formation across the syncytial nucleo-cytoplasmic domains, we performed photoactivation experiments in embryos defective in furrow formation. RhoGEF2 is a Rho-GTP exchange factor specifically needed for the formation of furrows in the syncytial embryo (Barmchi *et al*., 2005; Cao *et al*., 2008; Crest *et al*., 2012). Depletion of RhoGEF2 leads to shortened furrows and increase in RhoGEF2 is likely to increase active Myosin II and abolish furrow formation (Sherlekar and Rikhy, 2016; Zhang *et al*., 2018). We overexpressed RhoGEF2 by crossing flies containing mat-Gal4 and UASp-RhoGEF2. Embryos overexpressing RhoGEF2 showed short or missing furrows in metaphase (Figure 5A). The metaphase spindles did not show a significant change in these embryos as compared to controls (Figure 5A).

**Figure 5:**
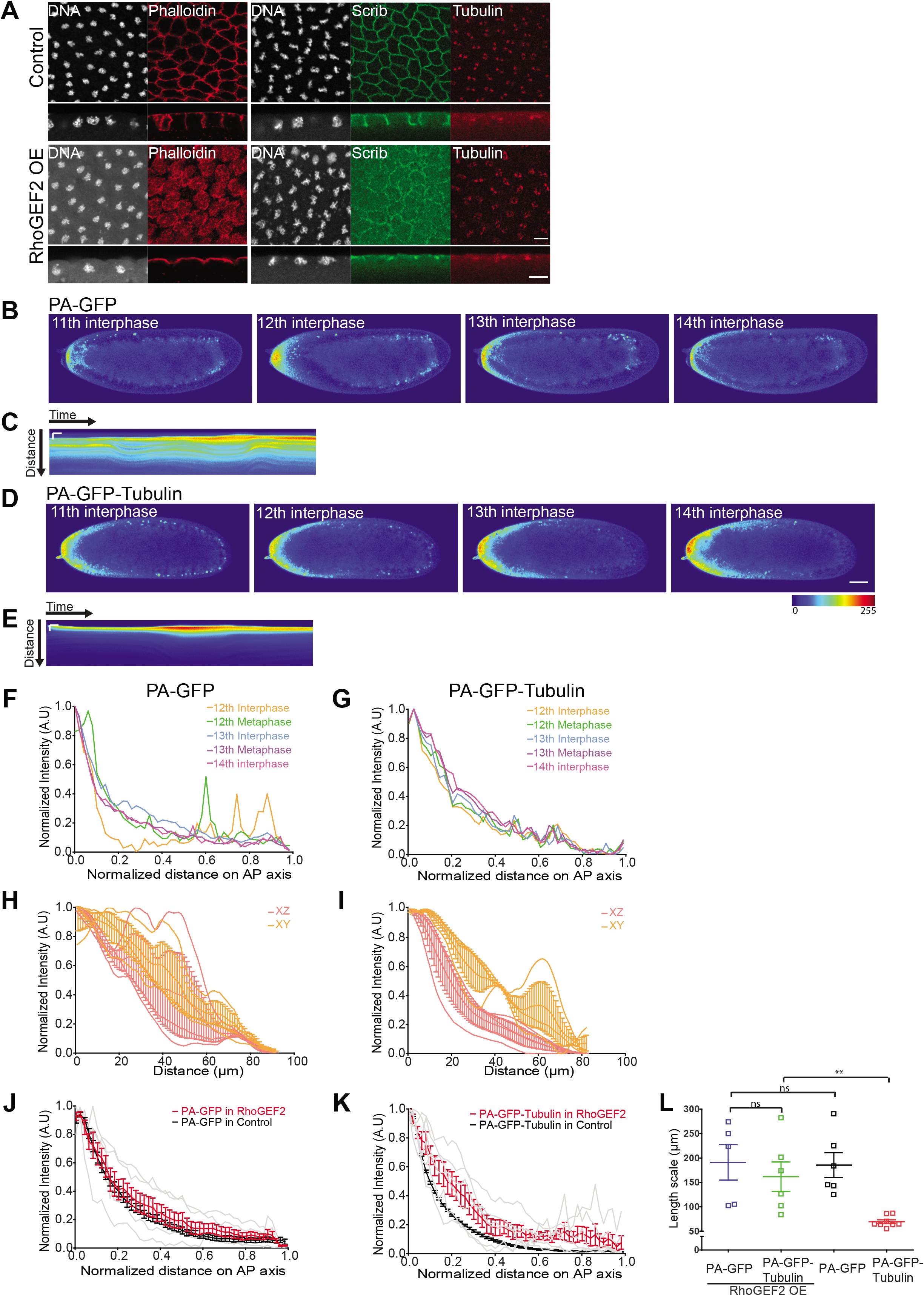
PA-GFP-Tubulin spreads to a greater extent in embryos containing RhoGEF2 overexpression. A. Embryos overexpressing RhoGEF2 show loss of metaphase furrows: Surface and sagittal views of fixed control or RhoGEF2 overexpressing embryos, stained with Tubulin (Red), Scribbled (Green) and DNA (Grey), show loss of furrows and no significant effect on metaphase spindles (100%, n=30 embryos). Scale bar=10μm. B-E. Images from syncytial cycles of an embryo co-expressing RhoGEF2 along with PA-GFP (B) or PA-GFP-Tubulin (D) which has been photoactivated at the anterior pole. Kymograph for PA-GFP (C) and PA-GFP-Tubulin (E) shows increase in cortical fluorescence with time while sometimes changing the extent to which the fluorescence is confined. Scale bar=50μm, 60s. F-G. Quantification of evolution of the photoactivation across nuclear cycles in embryos overexpressing RhoGEF2. A line profile was drawn in syncytial cycles is plotted for PA-GFP (F) and PA-GFP-Tubulin (G) for one embryo. Similar profiles were observed in multiple embryos (n=3 for each). H-I. Quantification of extent of photoactivation in XY vs XZ direction for PA-GFP (H) and PA-GFP-Tubulin (I) in embryos overexpressing RhoGEF2. The raw data is shown in a lighter color and the averaged data is shown in a darker color, error bars represent standard error on means (n=3 embryos for PA-GFP and PA-GFP-Tubulin each). J-K. Quantification of intensity profile of photoactivated probe as measured at the end of the experiment for PA-GFP (J) and PA-GFP-Tubulin (K) in embryos overexpressing RhoGEF2. The raw data is shown in a lighter color and the averaged data is shown in a darker color, error bars represent standard error on means (n=3 embryos for PA-GFP-Tubulin and PA-GFP each). The graph for photoactivation of PA-GFP and PA-GFP-Tubulin in control embryos is the same as that shown in Figure 3I and is repeated here for comparison. L. Scatter plot of the length scales extracted after fitting an exponential decay function to the intensity profiles seen in J and K. The values of length scales for PA-GFP and PA-GFP-Tubulin for anterior photoactivation in control embryos are repeated from Figure 3J. (n=5,3 for PA-GFP in RhoGEF2-OE, 6,3 for PA-GFP-Tubulin in RhoGEF2-OE. Two tailed Mann-Whitney non-parametric test, p value 0.53 for PA-GFP and PA-GFP-Tubulin in RhoGEF2-OE, 0.93 for PA-GFP/RhoGEF2-OE and PA-GFP/control, 0.002 for PA-GFP-Tubulin/RhoGEF2-OE and PA-GFP-Tubulin/control).

Next, we generated embryos expressing PA-GFP or PA-GFP-Tubulin along with RhoGEF2 overexpression. We performed continuous photoactivation at the anterior pole in a fixed area and followed the resultant gradient across time (Figure 5B,D, Movie S7,8). We found that similar to the control embryos (Figure 2), the gradients evolved over time (Figure 5B-G). Further, in spite of major contractions in the embryo yolk (Movie S7,8), the activated fluorescent molecules remained near the cortex and did not mix with the underlying inner yolk region of the embryo (Figure 5H,I). This was also evident from the kymographs which showed undulations in the cortical layer of fluorescence, yet maintaining a separation from the embryo’s inner yolk region (Figure 5C,E). The PA-GFP gradients did not change in the embryos over-expressing RhoGEF2 (Figure 5J). The PA-GFP-Tubulin gradient however changed significantly (Figure 5K) when compared to their respective gradients in control embryos (Figure 2). The length scales were extracted on fitting an exponential function to the concentration profile obtained. It was found that PA-GFP-Tubulin (161+30μm) gradient spread to a greater extent in embryos overexpressing RhoGEF2 as compared to control embryos. There was no significant difference in PA-GFP-Tubulin spread from PA-GFP (191+36μm) in RhoGEF2-OE embryos (Figure 5L). RhoGEF2 overexpression led to loss of plasma membrane furrows and loss of restriction of PA-GFP-Tubulin gradient in the syncytial *Drosophila* embryo.

### Anteriorly photoactivated PA-GFP-Tubulin gradient length scale increases in mutants of EB1

Microtubules emanate from the centrosome at the apical side and spread vertically downwards in the syncytial blastoderm embryo (Kellogg *et al*., 1988; Sullivan and Theurkauf, 1995). EB1 is present at the growing end of microtubules and its depletion is likely to disrupt the microtubule architecture (Rogers *et al*., 2002). We depleted embryos of EB1 by combining *eb1* RNAi to mat-Gal4 to disrupt microtubule organization. The microtubule staining was reduced in embryos expressing *eb1* RNAi expressing embryos. Also the plasma membrane levels for Scribbled were lowered (Figure 6A).

**Figure 6:**
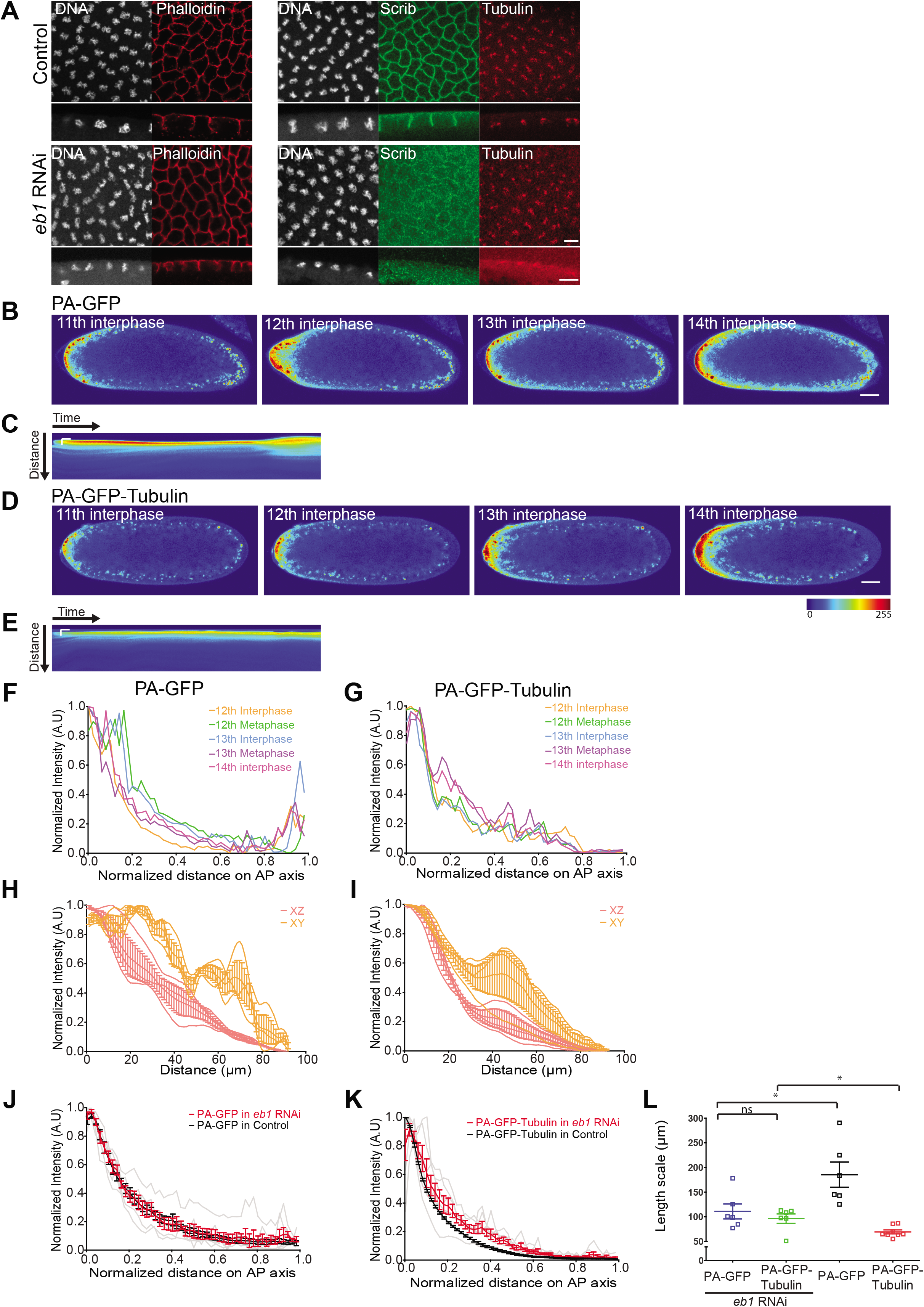
PA-GFP-Tubulin spreads to a greater extent in *eb1* mutant embryos. A. *eb1* RNAi expressing embryos show perturbed tubulin architecture: Surface and sagittal views of fixed control or *eb1* RNAi embryos, stained with Tubulin (Red), Scribbled (Green) and DNA (Grey), show perturbed spindles in metaphase (100%, n=25 embryos). Scale bar=10μm. B-E. Images from syncytial cycles of an *eb1* RNAi expressing embryo co-expressing PA-GFP (B) or PA-GFP-Tubulin (D) with photoactivation at the anterior pole. Kymograph for PA-GFP (B) and PA-GFP-Tubulin (E) shows increase in cortical fluorescence across time while sometimes changing the extent to which the fluorescence is confined. Scale bar=50μm, 60s. F-G. Quantification of evolution of photoactivated signal across nuclear cycles in *eb1* RNAi embryos. Graph depicts intensity change in PA-GFP (F) and PA-GFP-Tubulin (G) for one embryo. Similar profiles were observed in multiple embryos (n=3 for each). H-I. Quantification of the extent of photoactivated probe spread in XY vs XZ direction for PA-GFP (H) and PA-GFP-Tubulin (I) in *eb1* RNAi embryos. The raw data is shown in a lighter color and the averaged data is shown in a darker color, error bars represent standard error on means (n=3 embryos for PA-GFP and PA-GFP-Tubulin each). J-K. Quantification of intensity profile of photoactivated probe as measured at the end of the experiment for PA-GFP (J) and PA-GFP-Tubulin (K) in *eb1* RNAi embryos. The raw data is shown in a lighter color and the averaged data is shown in a darker color, error bars represent standard error on means (n=3 embryos for PA-GFP-Tubulin and PA-GFP each). The graph for photoactivation of PA-GFP and PA-GFP-Tubulin in control embryos is the same as that shown in Figure 3I and is repeated here for comparison. L. Scatter plot of length scales extracted after fitting an exponential decay function to the intensity profiles seen in J and K. The values of length scales for PA-GFP and PA-GFP-Tubulin for anterior photoactivation in control embryos are repeated from Figure 3J. (n=6,3 embryos for PA-GFP in *eb1* RNAi, 6,3 for PA-GFP-Tubulin in *eb1* RNAi. Two tailed Mann-Whitney non-parametric test, p value 0.81 for PA-GFP and PA-GFP-Tubulin in *eb1* RNAi, 0.02 for PA-GFP/*eb1* RNAi and PA-GFP/control, 0.04 for PA-GFP-Tubulin/*eb1* RNAi and PA-GFP-Tubulin/control).

We combined the *eb1* RNAi with flies expressing PA-GFP or PA-GFP-Tubulin and performed anterior photoactivation experiments in a fixed area (Figure 6B,D, Movie S9,10). We found that similar to the control embryos (Figure 2), the gradient evolved over time (Figure 6B-G). Similar to RhoGEF2 overexpression embryos, in spite of major contractions in the embryo yolk, the activated fluorescent molecules remained near the cortex (Figure 6C,E) and did not mix with the underlying yolk region of the embryo (Figure 6H,I). Length scales were extracted by fitting these gradients (Figure 6J,K) to an exponential function. We saw that the PA-GFP gradient did not change, while PA-GFP-Tubulin gradient in mutant embryos changed significantly. The length scale analysis showed that PA-GFP-Tubulin (96±9μm) in *eb1* RNAi spread more than control embryos. There was no significant difference between the PA-GFP-Tubulin length scale as compared to PA-GFP (111±15μm) in *eb1* RNAi embryos, even though the PA-GFP was more constrained than control embryos (Figure 6L). In summary, *eb1* mutant embryos showed a disrupted microtubule architecture and showed a loss of restriction of PA-GFP-Tubulin gradient in the syncytial *Drosophila* embryo.

## Discussion

In this study, we have examined the distribution and diffusion of cytoplasmic components of the *Drosophila* syncytial blastoderm embryo. We have used photoactivation of cytoplasmic PA-GFP to analyze its distribution and diffusion across nucleo-cytoplasmic domains of the syncytial *Drosophila* embryo and further compared it to PA-GFP-Tubulin, which is present in the cytoplasm and is also incorporated in microtubules. We find that the cytoplasmic components have an increased concentration at the cortex near the nucleo-cytoplasmic domains. Photoactivation of these components shows diffusion to a greater distance in the antero-posterior axis in the cortex as compared to the depth of the embryo. Also photoactivated cytoplasmic components diffuse less when generated at the center of the embryo as compared to the anterior. Diffusion is constrained by interaction with the cyto-architecture components of the syncytial blastoderm embryo (Figure 7).

**Figure 7:**
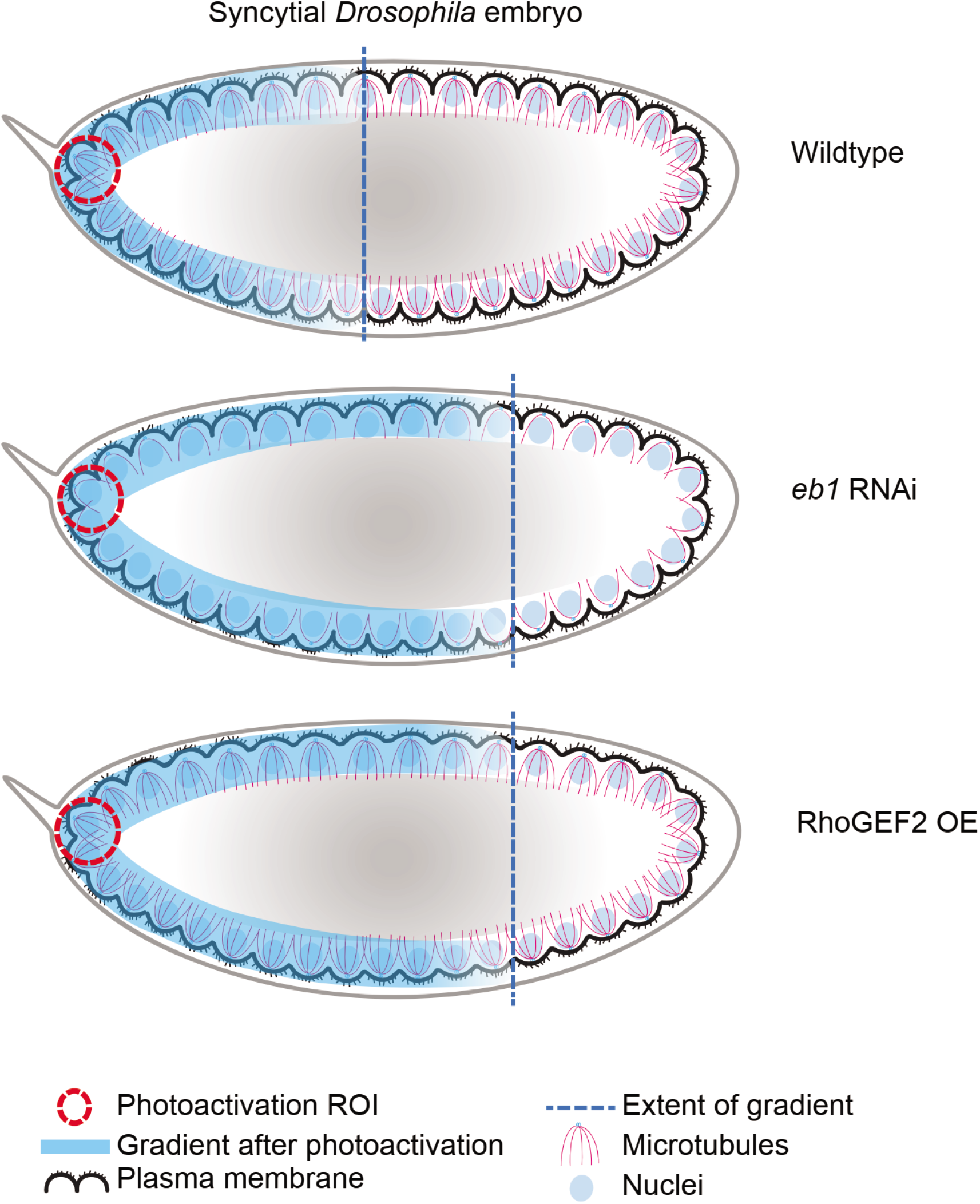
Model for regulation of gradient formation across the nucleo-cytoplasmic domains in the syncytial *Drosophila* embryo. Photoactivated PA-GFP-Tubulin and PA-GFP form a cortical gradient in the syncytial blastoderm embryo. RhoGEF2 overexpression causes loss of plasma membrane furrows and increased spread of the anteriorly induced PA-GFP-Tubulin gradient. EB1 loss causes perturbation of the microtubule cytoskeleton and increased spread of the anteriorly induced PA-GFP-Tubulin gradient.

### Photoactivation as a method to study regional differences in kinetics of gradient formation in the syncytial Drosophila embryo

The use of photoactivatable GFP molecules allows for the creation of localized ectopic gradients and enables us to follow their evolution in real time across the syncytial nuclear cycles. Photoactivation has been used previously to analyze the spread of morphogen gradients in similar contexts. Photoactivation of Dorsal-PA-GFP allowed an analysis of the extent of its spread in the dorsal versus the ventral side of the syncytial blastoderm embryo. Sequestration of Dorsal by signaling components and nuclear capture on the ventral side gave a more constrained gradient as compared to the dorsal side of the embryo (Carrell *et al*., 2017). In our study we used two photoactivatable proteins which are incorporated in all nucleo-cytoplasmic domains. This allows us to quantify the differences in their diffusion due to inherent differences in their association with cyto-architecture of the embryo. We found that PA-GFP and PA-GFP-Tubulin had smaller length scales when activated at the center as compared to the anterior of the syncytial blastoderm embryo. The restricted diffusion at the center of the embryo could be a result of a difference in relative crowding of nucleo-cytoplasmic domains in these two regions (Blankenship and Wieschaus, 2001; Rupprecht *et al*., 2017). An increase in the density of nucleocytoplasmic domains in the center could lead to greater sequestration of cytoplasmic components in general, leading to a smaller length scale. Alternatively this could also come about due to differences in protein degradation or sequestration machinery between these two regions. Whether the difference in density of nucleo-cytoplasmic domains also leads to change in viscosity in the two regions remains to be examined.

### Cytoplasm organization in cells

The cytoplasm of majority of living cells can be described as an inhomogeneous, multi-phasic medium. Images of different components when drawn to scale (Goodsell, 2013) clearly convey the fact that the cytoplasm is quite contrary to the earlier picture of a freely flowing medium. The cytoplasm can be likened to a complex medium comprising of physical constraints and constraints due to binding and crowding. Fluorescent dextran of various sizes when injected into cells partitions based on their size (Luby-Phelps, 2000). This further corroborates the fact that the space available for various cytoplasmic components is constrained depending on their size. The metabolic state can also change properties of the cytoplasm in the bacterial cell into either a glass-like or fluid-like state (Parry *et al*., 2014). Cytoplasmic distribution can change depending on the ability and strength of a cytoplasmic molecule to bind to other components. A modelling based study showed that binding to negative end directed dynein motors on the mitotic spindle was sufficient to partition the cytoplasm into two halves even without the presence of any membrane bound compartments (Chen *et al*., 2012).

Our finding that PA-GFP-Tubulin is more restricted in its spread as compared to PA-GFP suggests that cytoplasmic components having multiple interactors are more confined in their diffusion. For the syncytium, this property is beneficial, as components produced from a syncytial nucleus tend to remain near their parent nucleus, with no clear boundaries being present in the shared cytoplasm. This observation suggests that different components in a cell could be restricted by distinct mechanisms, some binding to microtubules, some to actin or some being sequestered in the nuclei or other organelles ultimately resulting in restricting their action in space and time. The diffusion of PA-GFP-Tubulin in our study increased on abrogation of the metaphase furrows and microtubule cytoskeleton in embryos over-expressing RhoGEF2 and *eb1* RNAi, it reached length scales similar to PA-GFP. This further suggested that binding and sequestration were responsible for PA-GFP-Tubulin restriction. Loss of plasma membrane furrows could also lead to disorganization of astral microtubules (Cao *et al*.,2010; Crest *et al*., 2012) in the periphery thereby increasing the effective diffusion of PA-GFP-Tubulin.

Further, the observation that cytoplasmic components are cortically enriched corroborates previously reported data about Bicoid movement in the cortex and its dependence on the actin and the microtubule cytoskeleton of syncytial blastoderm embryos (Cai *et al*., 2017). SEM images from cross sectioned *Drosophila* embryos also show the presence of similar biphasic compartments (Figard *et al*., 2013; Turner and Mahowald, 1976). Filamentous actin and non-muscle myosin are concentrated in the 3-4μm and 1-2μm region of the “ yolk-free” cytoplasm just beneath the plasma membrane of the embryo, respectively (Foe, Odell and Edgar, 1993). The cortical yolk-free cytoplasm increases in its depth as the syncytial cycles progress (Foe, Odell and Edgar, 1993). Our study is a further characterization of protein mobility in these phases, and we show that the cortical cytoplasm and yolk beneath it seem to form two separate phases, and do not mix in spite of major contractions of the embryo in mutant embryos. The size of the cortical cytoplasmic region as determined by the spread of cytoplasmic GFP in our study, is approximately 40μm, where the fluorescence intensity falls off sharply. This observation raises further questions about how these two phases are separate and the mechanisms that contribute to maintaining their integrity.

### Implications on morphogen diffusion

The observation about the presence of two separate phases of cortical cytoplasm and embryo yolk provides an interesting perspective to our current understanding of the morphogen gradients in the early embryo, namely, Bicoid, Dorsal and Torso. The Bicoid gradient has been extensively studied using the framework of the synthesis, diffusion and degradation (SDD) (Durrieu *et al*., 2018; Gregor *et al*., 2007; Grimm *et al*., 2010) and related models. However, a complete theoretical understanding of the mechanisms underlying the formation of the Bicoid gradient is still lacking. Our finding implicates a restriction of the effective volume in which Bicoid gradient develops and matures. It also raises the possibility that various cytoarchitectural components could impinge on its formation. For example, perturbations in furrows or cytoskeletal structures can change the effective concentration of morphogens in the cortical cytoplasm, leading to changes in the morphogen profiles, specifically for Bicoid.

There have also been various studies, implicating the size and shape of the mRNA source in Bicoid gradient formation (Fahmy *et al*., 2014; Little *et al*., 2011; Spirov *et al*., 2009). Photoactivation allows creation of different sized sources which can produce PA-GFP/PA-GFP-Tubulin or morphogen gradients at different rates and provides an opportunity to study the effect of the source on the gradient shape and dynamics.

The observation of distinct gradient length scales of PA-GFP-Tubulin versus PA-GFP points to another facet of morphogen gradient formation, namely decrease in the diffusivity of morphogens based on their interactions. FGF gradient is known to interact with Heparan sulfate proteoglycans which changes the effective diffusivity of the morphogen. The removal of these proteoglycans leads to an increase in the morphogen spread (Balasubramanian and Zhang, 2016). We can interpret the difference between the PA-GFP and PA-GFP-Tubulin profiles as being a consequence of increased binding of tubulin to the microtubule architecture. This leads to increase in its residence time by sequestration and thus a lower effective diffusion and consequently, a smaller length scale. It would be interesting to analyse the effect of removal of binding interactions for well-studied morphogen like Bicoid. It is notable that Dorsal gradient is known to be modulated depending on the presence or absence of a dimerizing GFP (Carrell *et al*., 2017).

Finally, our observation of difference in length scales between anterior versus centre photoactivation suggests a difference in cyto-architectural properties for different regions of the embryo. Our studies necessitate a systematic analysis of the impact of local architectural properties in the formation and maintenance of morphogen gradients.

### Materials and methods *Drosophila* stocks and crosses

*Drosophila* stocks were maintained in standard corn meal agar at 25°C. All crosses were set up at 25°C, except *eb1* RNAi (29°C). *mat-gal4-vp16; mat-gal4-vp16* (Girish Ratnaparkhi, IISER, Pune, India) was used to drive mCherry-alpha-TubulinA1B (mCherry-Tubulin) (#25774), PA-GFP (gift from Prof. Gerald M. Rubin, Janelia Research Campus, VA, USA), PA-GFP-alpha-Tubulin84B (PA-GFP-Tubulin) (#32076), UASp-RhoGEF2 (#9386) and *eb1* RNAi (#36599). GFP expressed under ubiquitin promoter (*ubi*-GFP, #1681) was imaged directly.

### Microscopy

1.5 hour old embryos were collected on sucrose agar plates, washed, dechorionated using 100% bleach, mounted on coverglass chambers (LabTek, Germany) in PBS (Mavrakis *et al*. 2008) and imaged on Plan-Apochromat 25x/0.8 Oil Immersion or Plan-Neofluar 40x/1.30 Oil objective on Zeiss LSM780 or LSM710 systems. PA-GFP and PA-GFP-Tubulin were photoactivated using the 405 nm diode laser using the bleach module on the LSM software. PA-GFP and PA-GFP-Tubulin thus produced was imaged using the 488 nm laser. ROI size was kept constant at 373μm^2^. Photoactivation iterations were kept constant at 10 iterations per frame with activation being performed after every frame. The photoactivation was carried out for 0.36s (10 iterations). 512 pixel × 512 pixel images were acquired after that with a scan speed of 1.97 seconds per frame. Mid sagittal sections were imaged. 8 bit images were acquired with mean line averaging of 2. The gain and laser power were adjusted to be cover the dynamic range of each fluorescent tag and care was taken to not reach 255 on the 8 bit scale. Pinhole was kept open at 180μm.

### Immunostaining

F1 flies were selected from Gal4 and mutant crosses were transferred to embryo collection cages (Genesee Scientific, CA, USA) with 2.5% sucrose agar supplemented with yeast paste. Embryos were washed, dechorionated using 100% bleach for 1 min, washed and fixed in heptane: 4% paraformaldehyde (1:1) in PBS (1.8 mM KH2PO4,137 mMNaCl, 2.7 mMKCl, 10 mM Na2HPO4) for 20 mins at room temperature. Embryos were devitellinized by vigorously shaking in heptane:methanol (1:1) for anti-Tubulin and Scribbled immunostaining. 2% Bovine Serum Albumin (BSA) in PBS with 0.3% Triton X-100 (PBST) was used for blocking. Following primary antibodies were diluted in the block solution: anti-Tubulin (Anti-mouse, Sigma-Aldrich, Bangalore, India,1:1000), anti-Scrib (Anti-Rabbit, Kind gift by Prof. Kenneth Prehoda, University of Oregon, OR, USA,1:1000). Fluorescently coupled secondary antibodies (Alexa Fluor 488, 568, 647 coupled anti-rabbit and anti-mouse, Molecular Probes, Bangalore, India) were used at 1:1000 dilution in PBST. Embryos were imaged using LD LCI Plan-Apochromat 25x/0.8 ImmKorr DIC M27 objective on the Zeiss LSM710/780.

### Image analysis

Segmented lines of 10 or 20 pixel (10 or 20μm width) were drawn across the cortex from the anterior to the posterior or centre to anterior/posterior of the embryo on the dorsal and the ventral side. Line profile measurements, containing embryo length vs intensity values were obtained using ImageJ. For XZ analysis, similar segmented lines were drawn for a distance of 90μm from the place of activation, in XY or XZ directions. The process was multiplexed using ImageJ macros. A MATLAB script was used to process the generated files. The script rescales the embryo length from 0 to 1 in the antero-posterior direction, subtracts the minimum intensity value, rescales it with the maximum and smoothens the intensity values using sliding window averaging.

### Sampling and Statistics

3 or more embryos as indicated in the corresponding figure legends were imaged and quantified for each experiment. Graphpad Prism 5.0 was used for Statistical analysis and plotting.

### Theory

#### Estimation of length scales from concentration profiles

The time evolution of the concentrations of the photo-activated molecules were analysed within the framework of the standard one-dimensional Synthesis-Diffusion-Degradation (SDD) model in a domain of length *L*(Crick, 1970),

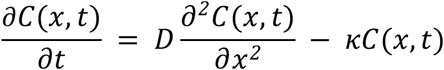

where, *C*(*x,t*) represents the concentration of the photo-activated species as position *x* at time *t, D* is the diffusion constant, and *κ*is the degradation rate. The mean lifetime of the molecule *τ*is the inverse of the degradation rate, *τ = 1/κ*. This equation is to be solved subject to the appropriate boundary conditions, accounting for the presence of a localised source of fluorescent molecules at the anterior pole of the embryo (or at the centre in the case of centre activation), and reflecting boundary conditions at the posterior pole,

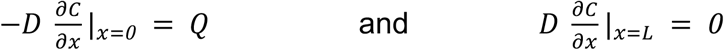

and the appropriate initial condition reflecting the absence of any photoactivated molecules for *t* ≤ 0, *C*(*x,t* ≤ 0) = 0.

At long enough times, the concentration profile evolves to a steady state (Figure 3C,D). The steady state solution of the SDD model for a semi-infinite domain is given by,

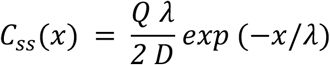

where the characteristic length-scale lambda is defined as, 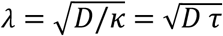. The semi-infinite assumption holds if the characteristic length-scale is much smaller than the size of the domain, *λ* ≪ *L*.

If the length scale is comparable to the system size, then the steady state solution depends on the length of the domain (size of embryo) and is given by,

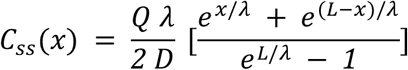

In order to ensure that the concentration profiles have reached a steady state, we plot the concentration versus time plots and the rate of change of concentration for both PA-GFP and PA-GFP-Tubulin. The time taken to reach the steady state depends on the position along the AP axis, and is smaller for locations closer to the anterior pole. We first show the results for PA-GFP-Tubulin (Figure 3F). As can be seen from the figures, the tubulin concentration reaches a steady state fairly quickly, justifying the assumption of the steady state for fitting the concentration profile. The time taken to reach the steady state can be determined by the time at which the derivative *dC/dt* approaches zero.

A similar analysis can be performed for PA-GFP (Figure 3E). The situation in this case is more complex, with the locations closer to the anterior pole having reached a steady state, while locations further away still evolving at the final time point of the experiments. The larger time taken to reach the steady state for PA-GFP can be understood from the fact that the length-scale for PA-GFP is much larger than PA-GFP-Tubulin and hence it takes a correspondingly larger time for the concentration profile as a whole to reach steady state. In this case, since the locations closer to the anterior pole have reached a steady state, we can fit the concentration profile in a localised region closer to the anterior pole.

The concentration profiles at the last time point are fitted by this steady state formula to obtain the characteristic length-scale *λ*. The fits are shown for PA-GFP(Figure 3G) and PA-GF-Tubulin (Figure 3H). This yields,

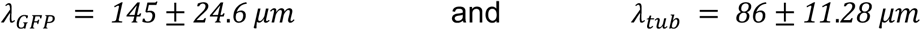

The PA-GFP spreads to a much larger distance from the anterior pole than Tubulin-PA-GFP.

#### Estimation of diffusion constant from concentration profiles

For the SDD model, the time taken to reach the steady state can be estimated theoretically (Berezhkovskii *et al*., 2010), and is given by,

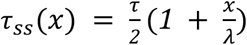

Where, *τ_ss_*(*x*)is the time taken to reach steady state at location *x*, and the mean lifetime of the molecule is denoted by *τ*, as before. The above formula also supports the notion that locations further away from the source at the anterior pole, take longer time to reach steady state. For distances much smaller than the characteristic length-scale, *x* ≪ *λ*, the above equation reduces to *τ_ss_*(*x*) = *τ*/2, and hence the mean lifetime can be read off from the concentration plots (Figure 3C,D) and its derivative plots (Figure 3E,F) by noting the time taken to reach steady state for both PA-GFP and PA-GFP-Tubulin for *x* = *11μm*(*x* ≪ *λ*). This yields,

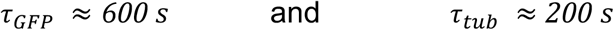

Combining the estimates of the length scale *λ* and the lifetime *τ*, we can then independently obtain an estimate of the diffusion constant,

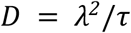

This gives,

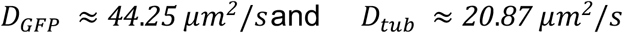

The estimation of the time taken to reach steady state makes certain assumptions. Firstly, the fluctuations in the concentration can be significantly high in certain embryos, which results in a large variation of the time estimate. Secondly, a characteristic feature of the time evolution of concentration profiles is that there is a sharp initial increase followed by a slow increase in the concentration. This suggests that there may be other biological processes beyond those described by the SDD model that affect the evolution of the concentration to the steady state. While estimating the diffusion coefficient, we neglect the slower variation and have chosen the onset of this slow increase as the steady state time.

## Abbreviations

(PA): Photoactivation
(RhoGEF2): Rho-GTP exchange factor 2

## Acknowledgements

Stocks obtained from the Bloomington *Drosophila* Stock Center (NIH P40OD018537) were used in this study. ST, BD, SS thank CSIR India for funding their fellowship. RR thanks DBT, DST and IISER Pune for funding the lab. BK thanks MHRD for fellowship. MM thanks DST Ramanujan Fellowship (13DST052) and IRCC, IIT Bombay for funding. AN acknowledges IRCC, IIT Bombay, India, and SERB, DST, India (Project No. ECR/2016/001967) for financial support.

## Supplementary Movies

S1: Cytoplasmic GFP: GFP expressed under the *ubi* promoter is imaged across the syncytial division cycles. Note that GFP enters the nuclei in interphase.

S2: mCherry-Tubulin: mCherry-Tubulin expressed with mat-Gal4 is imaged in the syncytial division cycles. Note mCherry-Tubulin incorporation into centrosome, spindle and cortical microtubules.

S3: PA-GFP anterior photoactivation: Region of interest at the anterior is photoactivated to create a source of PA-GFP. Note that PA-GFP enters the nuclei in interphase.

S4: PA-GFP-Tubulin anterior photoactivation: Region of interest at the anterior is photoactivated to create a source of PA-GFP-Tubulin. Note PA-GFP-Tubulin incorporation into centrosome, spindle and cortical microtubules.

S5: PA-GFP middle photoactivation: Region of interest in the middle of the embryo is photoactivated to create a source of PA-GFP.

S6: PA-GFP-Tubulin middle photoactivation: Region of interest in the middle of the embryo is photoactivated to create a source of PA-GFP-Tubulin.

S7: PA-GFP anterior photoactivation in RhoGEF2-OE embryos: Region of interest at the anterior is photoactivated to create a source of PA-GFP in RhoGEF2-OE embryos.

S8: PA-GFP-Tubulin anterior photoactivation in RhoGEF2 mutants: Region of interest at the anterior is photoactivated to create a source of PA-GFP-Tubulin in RhoGEF2-OE embryos.

S9: PA-GFP anterior photoactivation in *eb1* mutant embryos: Region of interest at the anterior is photoactivated to create a source of PA-GFP in *eb1* RNAi expressing embryos

S10: PA-GFP-Tubulin anterior photoactivation in EB1 mutants: Region of interest at the anterior is photoactivated to create a source of PA-GFP-Tubulin in *eb1* RNAi expressing embryos. Note the undulations caused by yolk contractions and that the cytoplasm remains peripheral, without mixing with the embryo yolk region.

All movies are in shown in 16 color intensity rainbow scale where Blue represents the lowest intensity and red represents the highest intensity. Scale bar=10μm or 50μm as mentioned.

